# Gut and Glomerular Barriers Determine Nanoplastic Fate and Systemic Impact

**DOI:** 10.64898/2026.01.28.699663

**Authors:** Melina Yarbakht, Mustafa Kocademir, George Sarau, Stefan Wirtz, Alexandra Ohs, Frank Schweda, Michelle Hinrichs, Mario Schiffer, Silke Christiansen, Janina Müller-Deile

## Abstract

Nanoplastics (NPs) are increasingly recognized as pervasive environmental toxicants, however, their interactions with gut and renal barriers, and the resulting systemic consequences remain poorly understood. Here, we studied the uptake of 50 nm polystyrene (PS) nanoparticles using a multi-scale approach integrating zebrafish models, isolated perfused mouse kidneys, and *in vitro* assays to delineate uptake and barrier-dependent organ distribution. In zebrafish larvae, PS-NPs were efficiently absorbed via the intestinal tract, as visualized by confocal and label-free stimulated Raman scattering (SRS) microscopy, leading to gut microbiota dysbiosis and systemic inflammatory responses. Despite widespread systemic dissemination, renal accumulation was minimal under physiological conditions, whereas both zebrafish and isolated perfused mouse kidneys exhibited substantial PS-NPs retention only when the glomerular filtration barrier was disrupted. *In vitro* glomerular endothelial cells and podocytes readily internalized PS-NPs without altering key glomerular identity markers, highlighting their intrinsic uptake capacity that is normally restricted *in vivo* by barrier integrity. Our findings establish the glomerular filtration barrier as a crucial gatekeeper that prevents renal nanoplastic deposition. Furthermore, we revealed a microbiota-mediated axis that may prime the kidney for the environmentally induced stressing in long term.

## Introduction

The ongoing rise in global plastic consumption has led to substantial amounts of plastic waste entering the environment ^1^. Plastics are inexpensive, durable, lightweight, and long-lasting materials. Microplastics (MPs) and nanoplastics (NPs) are generated through abrasion of tires and plastic waste or released via cosmetics, detergents, and textiles ^2, 3^. MPs and NPs are increasingly recognized as potential threats to human health. Environmental degradation, including ultraviolet radiation and biodegradation, can fragment plastic products into MPs (<5 μm) or even NPs (<100 nm) ^4^. These particles are now ubiquitous, with an adult human estimated to ingest 553 microplastic particles per day through air, food, or water, corresponding to 184 ng/capita/day ^5^. Polystyrene (PS), one of the most commonly studied plastics, is widely used in industry and everyday items such as toothbrushes ^2^.

Although MPs have been studied extensively, NPs likely pose a greater health risk due to their small size and potential tissue penetration. In aquatic organisms, particle size strongly influences biological impact; for example, 50 nm PS-NPs reduced survival in *Tigriopus japonicas* more than 0.5 µm particles ^6^. NPs can accumulate in multiple tissues, including brain, liver, muscle, and gut. In mice, PS-NPs have been detected in microglial cells, and size-dependent accumulation in liver, kidney, and intestine has been associated with oxidative stress, neurotoxicity, and disrupted energy balance ^7-10^. Emerging evidence implicates MPs and NPs in kidney injury. 2 µm PS-MPs induce mitochondrial dysfunction, ER stress, and autophagy in human proximal tubular epithelial cells ^11^, whereas PS-NPs trigger apoptosis, ROS production, and structural changes in mitochondria and endoplasmatic reticulum in a size- and dose-dependent manner ^9^.

Despite these insights, the mechanisms underlying NP uptake, distribution, and functional consequences at key organ barriers, such as the gut barrier and the glomerular filtration barrier, remain poorly understood. The gut barrier serves as the first line of defense against ingested particles, limiting their systemic translocation in healthy tissue. in a way that, the mucus layer and intact epithelial tight junctions trap the majority of particles, preventing entry into the bloodstream. Moreover, the gastrointestinal tract hosts a complex and dynamic microbial community that profoundly shapes host physiology, immune responses, and susceptibility to disease ^12, 13^. Exposure to NPs can compromise this barrier by downregulating tight junction proteins and increasing intestinal permeability ^14^, potentially facilitating systemic particle dissemination.

Following the entrance of particles in the circulation, they can reach the glomerular filtration barrier, which serves as the highly selective interface between blood and urine, restricting translocation based on size and charge under physiological conditions ^15^. Although occasional reports describe the presence of nanoparticles within renal tissue, the precise routes by which they access the kidney remain unclear ^11, 16-19^. Together, these observations suggest that the gut and glomerular filtration barriers are functionally interconnected in regulating nanoparticle biodistribution and that perturbations at one barrier may influence susceptibility at the other.

This uncertainty arises from two main factors, the high selectivity of glomerular filtration barrier which effectively prevents particles larger than a few nanometers from passing through ^15, 20^, and the insufficient *in vivo* evidence supporting alternative entry routes, such as transendothelial passage or tubular back-leak. Moreover, many studies showed that the renal nanoparticle localization with the use of fluorescent labeling or electron-dense proxies which can lead to artefactual aggregation and misidentification. This makes it experimentally challenging to confirm the true nanoparticle penetration into intact renal compartments. Recent evidence suggests that systemic pathways, including the gut–kidney axis influence renal responses to environmental toxicants ^21^. However, the contribution of plastic-derived nanoparticles to these interorgan interactions remains poorly resolved. Perturbations in the intestinal microbiota can modulate inflammation, oxidative stress, and metabolic homeostasis, thereby indirectly affecting kidney function ^22^ raising the possibility that microbiota-mediated effects may contribute to or amplify renal injury.

To address these knowledge gaps, we employed a multi-scale, integrative approach combining advanced imaging, microbiota profiling, bulk RNA sequencing, and physiological and functional assays to investigate how PS-NPs interact with both the gut barrier and the glomerular filtration barrier. Using *in vitro* studies with glomerular endothelial and podocyte cultures, *ex vivo* analyses in isolated perfused mouse kidneys, and *in vivo* experiments with zebrafish reporter lines, we bridge molecular, cellular, and organismal responses. Stimulated Raman scattering (SRS) microscopy provides a label-free, chemically specific method for visualizing nanoparticle distribution and retention in intact tissues by sensing their characteristic molecular vibrations. Recent advances have shown that hyperspectral SRS platforms can identify and image individual NPs below 100 nm with polymer-level specificity ^23^. This integrated strategy combining SRS imaging, zebrafish larval physiology, organ perfusion, microbiota analysis, gene expression analysis and cell culture provided an unprecedented resolution of NP delivery and enabled a rigorous examination of both direct and indirect cellular effects on the glomerular filtration barrier. These insights are crucial for evaluating the broader ecological and human health risks posed by MPs and NPs and for enhancing our understanding of how organism respond to exposure to environmental particles.

## Material and Methods

### PS-NPs characterization

Two types of carboxylated PS-NPs were used: fluorescent labeled 50 nm microspheres (Fluoresbrite®, excitation/emission 448/486 nm; Polysciences, Inc., Warrington, PA, USA) and non-fluorescent carboxylated microspheres (Polysciences, Inc., Warrington, PA, USA). Morphology was confirmed by scanning electron microscopy (SEM), while hydrodynamic size and zeta potential were measured using dynamic light scattering (DLS) (Malvern Panalytical, Malvern, UK). We used a concentration of 50 µg/mL PS-NPs to ensure the robust and measurable uptake of NPs in zebrafish kidney tissues, and to allow the precise tracking of particles. This dose is consistent with standard practices in nanotoxicology and zebrafish research ^24, 25^. Although the selected dose does not replicate realistic environmental exposures, it is crucial for thorough mechanistic evaluation and ensures reproducible, interpretable results related to nanoparticle uptake, biodistribution, and interaction with renal tissues.

### In vitro cell experiments

Conditionally immortalized human podocytes and glomerular endothelial cells (GECs) were used to study NPs uptake. Podocytes were cultured in RPMI 1640 medium (Gibco, Thermo Fisher Scientific, Waltham, MA, USA) supplemented with 10% fetal calf serum (PAN Biotech, Aidenbach, Germany), 1% penicillin–streptomycin (Sigma-Aldrich, St. Louis, MO, USA), and insulin–transferrin–selenium supplement (Gibco, Thermo Fisher Scientific, Waltham, MA, USA). GECs were cultured in Endothelial Cell Basal Medium MV2 (PromoCell, Heidelberg, Germany) with growth supplement (Sigma-Aldrich, St. Louis, MO, USA). Cells were maintained at 37 °C. Cells were treated with 25 µg/mL or 50 µg/mL green fluorescent PS-NPs for 48 h, washed with PBS, and fixed with 4% paraformaldehyde (PFA; Sigma-Aldrich, St. Louis, MO, USA). Lysosomes were labeled using LysoTracker**®** (Thermo Fisher Scientific, Waltham, MA, USA), and cytoskeleton filaments were stained with phalloidin (Thermo Fisher Scientific, Waltham, MA, USA). 3D reconstruction and NP quantification were performed using z-stacks acquired on a Leica TCS SP8 confocal microscope (Leica Microsystems, Wetzlar, Germany) and processed in ImageJ2 (National Institutes of Health, USA) with the Particle_in_Cell_3D macro ^26^.

### Zebrafish husbandry and experimental design

Zebrafish (*Danio rerio*) larvae with the nacre background were maintained under standard laboratory conditions. We applied three transgenic lines including *Tg(l-fabp:VDBP-eGFP)* for indirect assessment of loss of circulating proteins ^27^, *pod:NTR-mCherry* for podocyte-specific injury ^28^ and Tg(lyz:DsRed2) zebrafish labeling a subset of myeloid cells, including macrophages and some granulocytes.

Loss of circulating protein model: Tg(l-fabp:VDBP-eGFP) larvae were exposed to non-labeled PS-NPs at 3 days post fertilization (dpf) for 48 h. GFP fluorescence in retinal vessels was quantified using ImageJ (Version 1.48; National Institutes of Health, USA). Loss of fluorescence indicated proteinuria or vasculature leakage ^27^.

Podocyte injury model: At 3 dpf, double transgenic larvae were screened for mCherry-positive glomeruli and treated with metronidazole (80 µM or 1 mM; Sigma-Aldrich, St. Louis, MO, USA) in 0.1% DMSO/E3 medium for 48h. Control larvae was treated only with 0.1% DMSO. Loss of plasma protein assessment was done at 5 dpf to evaluate the effect of metronidazole. In order to track the uptake of NPs, in a parallel experiment, following the treatment of pod:NTR-mCherry larvae, zebrafish were treated with GFP-labeled PS-NPs at 3 dpf for 48 h. Furthermore, this experiment was done with a higher concentration of Metronidazole (1 mM) to assay the effect of it on the uptake of GFP-labeled PS-NPs.

Immune cell recruitment model: Tg(lyz:DsRed2) zebrafish ^29^ were exposed to GFP labeled PS-NPs at 3 dpf for 48 h. Whole zebrafish larvae were imaged using ACQUIFER imaging machine (Bruker, Heidelberg, Germany).

### Microplastic detection in zebrafish using SRS and fluorescence microscopy

Microplastic localization in zebrafish cross-sections was assessed using a Leica SP8 CARS confocal system with an integrated Stimulated Raman Scattering (SRS) module (Leica Microsystems, Wetzlar, Germany). The PicoEmerald S OPO laser generated picosecond pulses (fixed Stokes 1031.306 nm, tunable pump 720–980 nm) synchronized at 80 MHz. A 20× water-immersion objective (HC PL IRAPO, NA = 0.75) was used for imaging, with forward-scattered signals collected via a 0.9 NA dry condenser. Pump beam modulation at 20 MHz was detected using a lock-in amplifier (Zurich Instruments, Zurich, Switzerland). Laser powers were 0.3 W (Stokes) and 0.15 W (pump), with ∼30% applied during imaging.

SRS hyperspectral data were collected in the fingerprint region (980–1700 cm^−1^, 75 points) and C–H stretching region (2800–3200 cm^−1^, 32 points) at a single z-position. Images (512 × 512 pixels) were acquired at 15.38 µs dwell time, 100 Hz scan speed, and 2× line averaging. Regions of interest were pre-identified using fluorescence signals arising from labeled PS-NPs and endogenous tissue autofluorescence, enabling precise spatial correlation with SRS imaging.

Fluorescence microscopy was performed using the same system (excitation 481 nm, emission 491–541 nm), with HyD X1 detectors and ∼2 mW laser excitation power. This integrated approach allows concurrent acquisition of label-free chemical vibrational spectroscopic data, and fluorescence-based particle tracking, providing complementary chemical and spatial information within intact zebrafish tissue.

### Isolated perfused kidney

The mouse isolated perfused kidney was used to evaluate the detectability of the infused PS-NPs in the tissue samples. The procedure was performed with mice (C57BL/6 background) as kidney donors in two different groups with and without puromycin (1 mg/mL, 2 h), and 6.7 mg of PS-NPs which was added to the perfusate for 30 min. The isolated perfused kidney model was published before ^30, 31^. In brief, the mice were sacrificed by cervical dislocation, the abdominal cavity was opened and a metal perfusion cannula was inserted into the abdominal aorta and advanced to the origin of the right renal artery. The perfusion was started and the aorta was ligated proximal to the right renal artery. The right kidney was excised, placed in a thermostated moistering chamber, and perfused at a constant pressure of 100 mmHg using a modified Krebs-Henseleit solution including albumin, physiological amino acids and glucose ^30, 31^. After the infusion period with PS-NPs with or without puromycin, the kidneys were perfused with 4% paraformaldehyde (PFA, pH 7.4) for tissue fixation. Fixed kidney tissues were subsequently processed for stimulated Raman scattering (SRS) microscopy using the same spectral acquisition parameters described for zebrafish samples, enabling direct, label-free detection of PS-NPs within renal compartments.

### RNA isolation, cDNA transcription, and qPCR

RNA was isolated from whole cell lysates using the ReliaPrep™ RNA Miniprep System (Promega, Madison, WI, USA), following the manufacturer’s protocol. To synthesize cDNA, 1 μg of RNA was combined with the following reagents: 5 × RT buffer and M-MLV RT 50.000 U (Promega), dNTP Mix (Promega), random hexamer primer (ThermoFisher Scientific), and RiboLock (Thermo Fisher Scientific). Reverse transcription was done at 25 °C for 5 min, 40 °C for 60 min, and 70 °C for 10 min. Sybr green-based real-time PCR (Maxima SYBR Green/ROX qPCR Master Mix, Thermo Fisher Scientific) was performed as follows: 10 min at 95 °C followed by 40 cycles of 15 s at 95 °C and 1 min at 60 °C, followed by 15 s at 95 °C, 1 min at 60 °C and 15 s at 95 °C. Individual samples were run in triplicate. Data was analyzed by using the ΔΔCt method. Primer sequences were listed in **supplementary table 1**.

### RNA-seq

RNA sequencing was performed on zebrafish larvae (*Danio rerio)* using duplicate replicates of larvae exposed to PS-NPs and control larvae. RNA quality was assessed with a Agilent Tapestation system. Libraries were prepared according to Novogene’s in-house protocol and sequenced with 150 bp paired-end reads on an Illumina NovaSeq X (Illumina, USA), generating an average of 20 million reads per sample. Reads were quality-checked using FastQC (v0.12.1), trimmed with FASTP (v0.23.4) and aligned to the *Danio rerio* reference genome (GRCz11) with STAR (v2.7.10a). Salmon (v1.10.1) was used to generate count tables. Data normalization, exploration and differential abundance analysis were conducted in R (v4.1.1) using DESeq2 (v1.50.0). Principal Component Analysis (PCA) was performed using the plotPCA function and visualized with ggplot2 (v3.3.6). DEGs were visualized in volcano plots using ggplot2. Clusterprofiler (v4.18.0) was used for gene ontology analysis.

### Gene Set Enrichment Analysis (GSEA)

GSEA was performed with 100,000 permutations using a weighted enrichment score and pre-ranked genes ^32ref^. All genes with Entrez IDs were included, ranked by log2 fold-change values. Analyses employed Gene Ontology and Hallmark gene sets from MsigDB (v7.5.1) for *Danio rerio*. Functional enrichment was conducted using the *fgsea* package (v1.20.0), applying the Multilevel function with minGSSize = 5 and maxGSSize = 800. Adjusted p-values were calculated using the Benjamini-Hochberg method. Significant pathways (adjusted p < 0.05) were visualized using *ggpubr* (v0.6.0), and gene annotations were added with org.Dr.eg.db (v3.14.0).

### Intestinal microbiota analysis

To assess the effects of PS-NPs on the zebrafish microbiome, 3 dpf larvae were exposed to 50 µg/mL GFP-labeled PS-NPs for 48h. Larvae were anesthetized using 0.04% tricaine (Sigma-Aldrich, St. Louis, MO, USA), euthanized on ice, and intestines were dissected. DNA was extracted from 10 control larvae and 11 PS-NP treated larvae using the ZymoBIOMICS DNA Microprep Kit (Zymo Research, Irvine, CA, USA) according to the manufacturer’s instructions. Library generation, sequencing, and bioinformatic analysis described below were performed in the Core Unit Microbiome Analysis Center (MACE) of Erlangen.

### 16S-metabarcoding based microbiome analysis

Genomic 16S rRNA V4 regions were amplified from 10 ng of genomic DNA using the prokaryotic primer pair 515F (5′-GTGYCAGCMGCCGCGGTAA−3′) and 806R (5′-GGACTACNVGGGTWTCTAAT−3′) containing barcoded forward primers (https://earthmicrobiome.org/protocols-and-standards/16s/). PCR amplification was performed with NEBNext Q5 Hot Start Hifi Master Mix for 25 cycles. Amplicons were purified with AMPure XP beads, pooled equimolarly, and sequenced (2 × 151 bp paired-end) on an Illumina MiSeq platform. Raw FASTQ files were processed in QIIME2 (v2025.4) using DADA2 for quality control, dereplication, and generation of amplicon sequence variant (ASV) tables. Taxonomic classification was performed against a classifier trained against the SILVA 138.2 SSU rRNA database (99% similarity threshold). ASV and taxonomy tables were imported into R (v4.5) as a *phyloseq* object. Alpha and beta diversity analyses were conducted using the *vegan* (2.6.5) package after log-transformation and rarefaction to the smallest library size. Graphical representations were generated with *ggplot2*.

### Statistical analysis of bioinformatics data

To assess gut microbial community structure, Fisher’s alpha, Shannon indices and inverse Simpson diversity were calculated. For statistical comparison of alpha diversity between groups, the Wilcoxon rank-sum test was used. Beta diversity was evaluated using Bray-Curtis and Jaccard distance matrices and visualized via Principal Coordinate Analysis (PCoA). Permutational multivariate analysis of variance (PERMANOVA) was conducted using the *adonis2* function of *vegan* v2.6.5 with 9999 permutations. Unpaired t-tests were used for comparison between two groups. Statistical significance was defined as p <0.05. Imaging was performed under standardized conditions for intensity and background. Functional potential of the intestinal microbiota was inferred with PICRUSt2 based on Kyoto Encyclopedia of Genes and Genomes (KEGG) pathway annotations. Unpaired t-tests were used for comparison between two groups. Statistical significance was defined as p <0.05. Imaging was performed under standardized conditions for intensity and background.

## Results and Discussion

### PS-NPs have distinct SRS spectroscopy

Scanning electron microscopy (SEM) and dynamic light scattering (DLS) confirmed that the PS-NPs we used for our study were uniformly sized, spherical particles with an average diameter of ∼50 nm (**Fig. 1A, B**). To chemically verify the identity of the nanoparticles, we performed stimulated Raman scattering (SRS) spectroscopy, which detects molecules through their characteristic vibrational patterns. In SRS, specific molecular bonds absorb two synchronized laser beams only when their energy difference matches the natural vibration of that bond, producing a highly selective chemically signal that can be plotted as a spectrum ^13^. The PS-NPs exhibited distinct peaks at 1008 cm^−1^ and 1605 cm^−1^ within the so-called fingerprint region, a spectral range where molecules show highly specific vibrational features here corresponding to aromatic ring breathing and C=C stretching of polystyrene. In addition, a peak at 3068 cm^−1^ appeared in the CH-stretching region, a higher-energy range dominated by vibrations of carbon–hydrogen bonds, further confirming the presence of aromatic C–H groups characteristic of polystyrene (^23, 33^; **Fig. 1C**).

**Figure 1:**
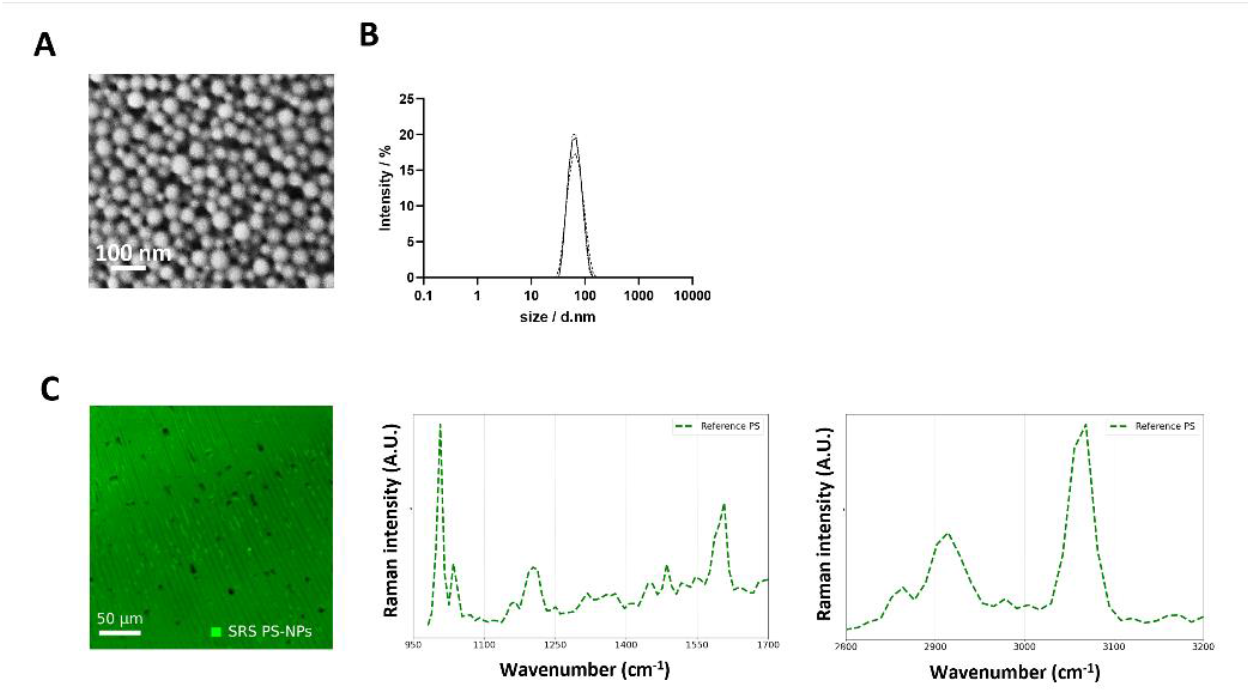
PS-NP characterization. A: Scanning electron microscopy of monodispersed spheric PS-NPs. Scale bar: 00 nm. B: Dynamic light scattering of PS-NPs with an average size of 50 nm. C: SRS image of PS-NPs acquired at the 1008 cm^−1^, together with corresponding SRS spectra showing characteristic peaks in the fingerprint region (1008 and 1605 cm^−1^) and in the C–H stretching region (3068 cm^−1^). Scale bar: 50 µm.

### Stimulated Raman scattering reveals spatial localization of PS-NPs in zebrafish larvae

To evaluate the *in vivo* distribution of plastic nanoparticles, 3-day-old zebrafish larvae were exposed to green fluorescent PS-NPs in fish water for 48 h and cryosections were analyzed at 5 dpf using combined fluorescence and SRS imaging. The zebrafish (*Danio rerio*) is a well-established model for studying the effects of environmental toxins, due to its optical transparency, rapid development, and genetic tractability. Zebrafish larvae develop a fully functional pronephric kidney and intestinal system, including a gut microbiota that, although simpler, shares key compositional features with that of humans ^34^. Zebrafish have also been extensively used to model glomerular injury ^35, 36^ as well as exposure to environmental particles ^37^. Zebrafish pronephros lacks a mature peritubular vasculature during early development. As a result, researchers can study the nanoparticle access to the kidney exclusively through the glomerular filtration barrier ^38^.

Unlike fluorescence microscopy, SRS offers label-free chemical contrast by capturing the intrinsic vibrational signals of both synthetic particles and biological molecules **(Fig. 2A)**. Combination of fluorescence channel for the tissue (405 nm, blue channel), and SRS channel for PS-NPs reveal the accumulation of nanoparticles relative to the surrounding intestinal tissue (green channel). This multimodal approach allows clear distinction between PS-NPs and the lipid- and protein-rich compartments of the tissue. Distinct PS-associated Raman peaks (1008, 1605, and 3068 cm^−1^) were detected within the zebrafish intestinal epithelium (**Fig. 2B, C**) following PS-NP exposure via the water. Tissue-rich regions exhibited a dominant peak at 1470–1475 cm^−1^ (CH_2_/CH_3_ bending), enabling spatial localization of PS-NPs within the lipid/protein matrix. The lipid-associated CH_2_ stretching vibration at 2850 cm^−1^ served as an additional reference to delineate tissue architecture and lipid-rich areas, providing further spatial context for particle localization within the complex tissue matrix.

**Figure 2:**
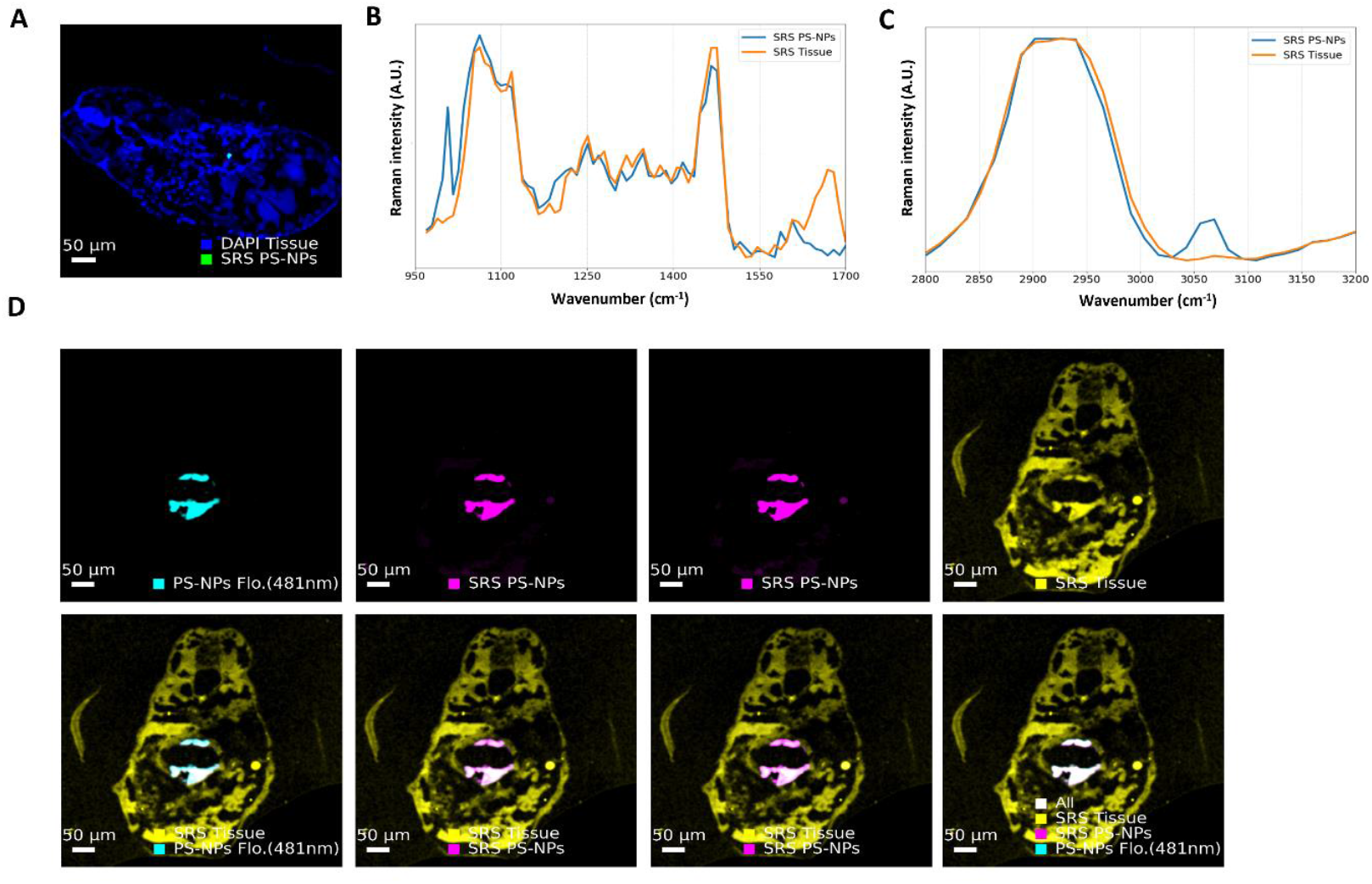
Stimulated Raman scattering (SRS) can detect PS-NPs in zebrafish tissue. A: Combined tissue fluorescence (405 nm, blue) and SRS imaging of zebrafish larval sections after 48 h exposure to PS-NPs (50 µg/L). Autofluorescence delineates the intestinal tissue architecture, whereas SRS at 1008 cm^−1^ identifies PS-NP-rich regions. Green and blue spectral traces correspond to PS-NPs and surrounding tissue background, respectively. Scale bar: 50 µm. B: SRS spectra from 1700–980 cm^−1^ showing the fingerprint region of PS-NP-rich areas (blue) and surrounding tissue (orange). The peak at 1008 cm^−1^ identifies the aromatic ring-breathing mode of PS-NPs, whereas the 1667 cm^−1^ band reflects endogenous protein and lipid signals. C: SRS spectra from the C–H stretching region showing PS-NP-rich areas (blue) and tissue (orange). The 3068 cm^−1^ peak corresponds to the aromatic C–H stretch of PS-NPs, while the 2850 cm^−1^ band indicates lipid-rich tissue. D: Fluorescence (481 nm) and SRS images acquired at 1008 cm^−1^ and 3068 cm^−1^ (PS-NPs) and at 2850 cm^−1^ (tissue lipids). The lower panel shows merged combinations of these channels, illustrating the spatial distribution of fluorescence-labelled PS-NPs relative to lipid and protein structures. Scale bar: 50 µm.

Complementary fluorescence imaging under identical optical conditions confirmed colocalization of green fluorescent PS-NP fluorescence within the intestinal lumen (**Fig. 2A, D**). This combined fluorescence-SRS visualization in Fig. 2D verifies that the Raman-identified particles correspond to the same PS-NP population detected by fluorescence, providing dual-modal confirmation of nanoparticle uptake. Larval zebrafish rely on yolk for nutrients until digestive tract opens and enzyme secretion begins at 5 dpf ^39-41^. Therefore, we hypothesize that during the period of 3-5 dpf, the uptake of NPs in the intestine occurs passively rather than through active feeding.

These data confirm the *in vivo* presence of PS-NPs and demonstrate the label-free chemical specificity of SRS in differentiating synthetic particles from surrounding biological components, facilitating precise spatial mapping of nanoparticle accumulation.

### PS-NPs induce edema without accumulating in the kidney in zebrafish larvae

Exposure of zebrafish larvae to labeled PS-NPs caused edema and led to marked NP deposition beneath the yolk, corresponding to the developing gut (**Fig. 3A**). We applied a transgenic zebrafish line expressing a green fluorescent circulating vitamin D-binding protein (VDBP-GFP), which allows indirect assessment for loss of proteins from the circulation as a cause for edema. A decrease in this albumin-sized plasma protein can easily be visualized in the retinal plexus ^42-44^. Notably, no significant reduction in circulating plasma protein was detectable after PS-NP exposure (**Fig. 3B**). SRS imaging of Tg(pod:NTR-mCherry) zebrafish, which labels podocytes in the glomerulus in red, confirmed the absence of PS-NPs in kidney tissue (**Fig. 3C**), consistent with the size- and charge-selective properties of the glomerular filtration barrier. Together, these results indicate that, despite systemic exposure and gut-associated retention, PS-NPs are effectively excluded from the kidney under physiological conditions, highlighting the glomerular barrier as a robust and selective defense against nanoparticle renal accumulation.

**Figure 3:**
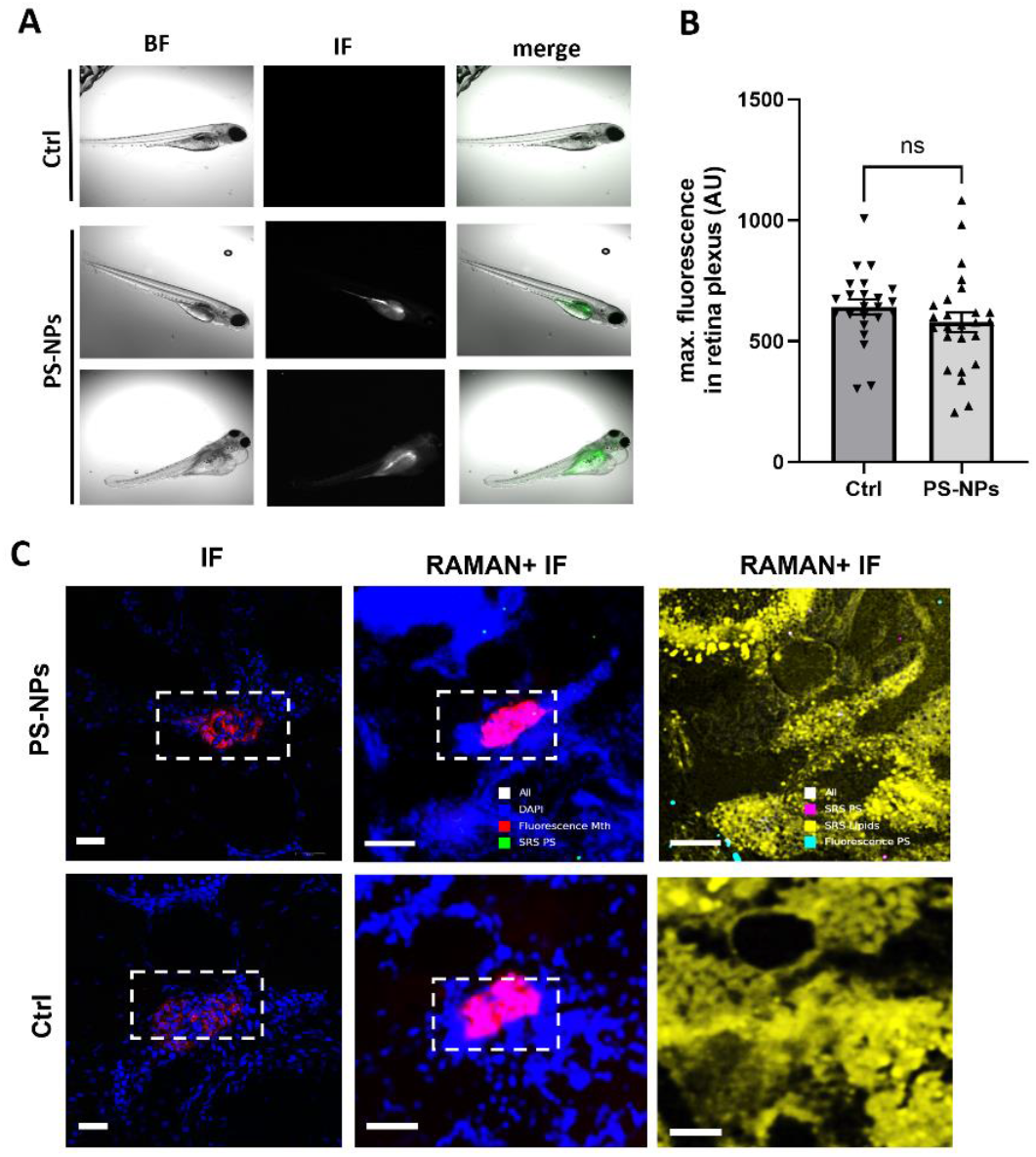
Exposure of zebrafish larvae to PS-NP causes edema with no evident accumulation of particles in the kidney. A: Phenotype pictures of zebrafish larvae of control group (Ctrl) versus larvae exposed to PS-NPs. Pictures were taken at 5 dpf after exposure to 50 µg/mL PS-NPs for 48 h. B: Proteinuria assay in Tg(l-fabp:VDBP:eGFP) zebrafish exposed to PS-NPs within the fish water or fish water only (Ctrl). The graph gives the maximum fluorescent intensity of the green fluorescent *VDBP* plasma protein measured in the retinal plexus of fish. Reduced signal is a sign for proteinuria. Assay was performed at 5 dpf after exposure to 50 µg/mL NPs for 48 h. n = 23 single larvae for PS-NP treatment and 20 control (Ctrl) single larvae per group. Differences were calculated between means ± SEM. N.s.: not significant. The graph represents one of five independent experiments conducted with comparable numbers of fish, all of which yielded consistent results with no significant differences observed. C: Multichannel SRS and fluorescence imaging of zebrafish larval sections exposed to green fluorescent 50 nm PS-NPs within the fish water or fish water only (Ctrl). Tissue-rich regions show a dominant peak around 1475 cm^−1^ (CH_2_/CH_3_ bending) and a reference peak at 1667 cm^−1^ (C=C stretching / amide I). PS-NPs were detected at 1008, 1605, and 3068 cm^−1^, and fluorescence of labeled PS-NPs was measured at 481 nm. The overlay image in the lower-right panel shows combined SRS and fluorescence channels. Scale bar: 50 µm.

The observed edema is likely caused by multiple factors. Zebrafish larvae depend on their skin for osmoregulation which exposes a large surface area to NPs. Dysfunction in skin tissue due to NP interactions may lead to fluid retention, independent of kidney injury. Thus, while gut barrier perturbation and microbial changes may also play a role, edema cannot be solely attributed to kidney dysfunction. This highlights the importance of considering the contribution of multi-organs in larval models.

Interestingly, our observation that PS-NPs did not accumulate in the zebrafish kidney or reach tubular cells are in contrasts with previous rodent studies which report detectable particles in those organs ^16^. However, the number of NPs detectable in this study was extremely low with only some detectable particles.

In mammals, some nanoparticles may reach the renal parenchyma not only via glomerular filtration but also through the extensive network of peritubular capillaries that surround the nephron. In contrast, zebrafish larvae lack a fully developed renal microvasculature at the larval stage and do not possess peritubular capillaries. Consequently, this model clearly lacks systemic distribution routes that are essential for effective secondary capillary uptake. Thus, the restricted renal localization of PS-NPs in zebrafish probably reflects a genuine physiological limitation of the larval kidney architecture, rather than a fundamental discrepancy in particle behavior. Importantly, the glomerular filtration barrier in zebrafish larvae is both structurally and functionally homologous to that of mammals, consisting of fenestrated endothelial cells, a trilaminar glomerular basement membrane, and interdigitating podocyte foot processes. Therefore, our data exclude glomerular filtration as an entry route for PS-NPs under baseline conditions, supporting the conclusion that direct passage of 50 nm particles across the filtration barrier is prevented by conserved size- and charge-selective properties.

### PS-NPs accumulate in glomeruli following podocyte injury in living zebrafish

To investigate whether glomerular barrier integrity controls the distribution of 50 nm green fluorescent PS-NPs in *in vivo* situation, we employed a zebrafish injury model, which allows real-time observation of renal function and nanoparticle interactions in a whole-organism context. Unlike the isolated perfused mouse kidney, which provides a controlled environment to study particle-kidney interactions independent of systemic and intestinal barriers, the zebrafish model incorporates the full physiological milieu, including circulation, glomerular filtration dynamics, and systemic responses. Therefore, we utilized the podocyte-specific ablation model Tg(pod:NTR-mCherry), in which podocytes express the bacterial enzyme nitroreductase (NTR) fused to mCherry under control of the podocin promoter. Upon treatment with metronidazole (MTZ), NTR converts this otherwise inert prodrug into a cytotoxic metabolite, leading to selective ablation of NTR-expressing podocytes. This model allows the controlled, temporally precise induction of podocyte injury and disruption of the glomerular filtration barrier **(Fig. 4A)** ^45^. Tg(pod:NTR-mCherry) zebrafish were crossed with our proteinuria reporter line (Tg(l-fabp:VDBP-eGFP) larvae) to induce podocyte damage and assess loss of circulating plasma protein in parallel. In contrast to treatment with 50 nm unlabeled PS-NPs, MTZ treatment (80 µM) for 48 h induced severe proteinuria, evidenced by the loss of green fluorescence of circulating protein **(Fig. 4B)**. Notably, combined MTZ and PS-NP treatment did not exacerbate plasma protein loss compared to MTZ alone, suggesting that PS-NPs do not further compromise podocyte function or glomerular filtration under conditions of pre-existing barrier injury **(Fig. 4B)**.

**Figure 4:**
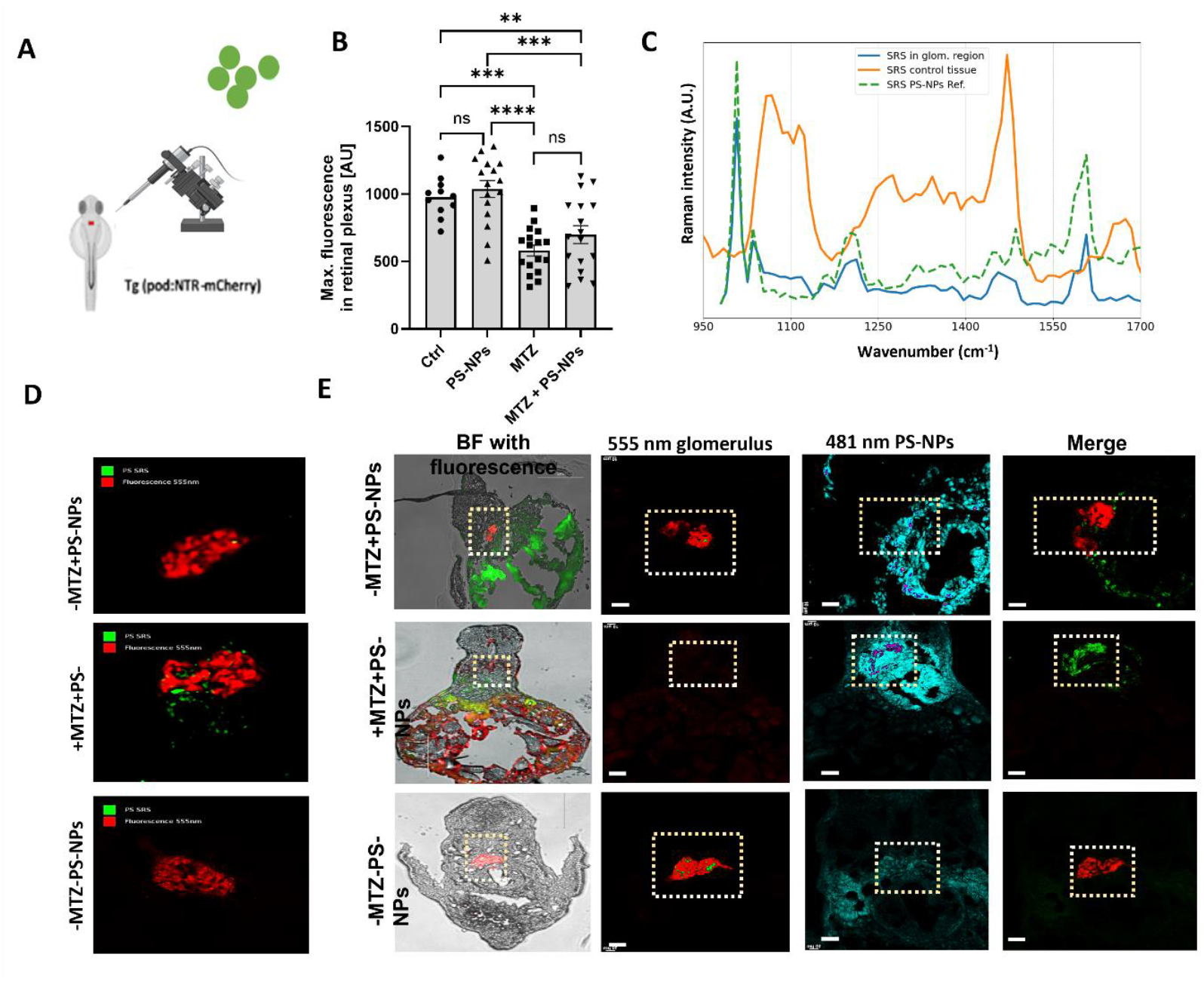
PS-NPs accumulate in glomerular regions after glomerular damage in vivo. A: Schematic illustration of 50 nm PS-NP injection into the Tg(pod:NTR-mCherry) zebrafish line. B: Proteinuria assay in Tg(l-fabp:VDBP-eGFP);Tg(pod:NTR-mCherry) zebrafish exposed to PS-NPs, metronidazole (MTZ), or the combination (PS-NPs +MTZ). Maximum VDBP-eGFP fluorescence intensity was measured in the retinal plexus at 5 dpf after 24 h exposure to 80 µM MTZ. n = 11 (Ctrl), 17 (MTZ), 17 (PS-NPs), 16 (PS-NPs +MTZ). Data represent mean ± SEM. n.s., not significant; **p < 0.01; ***p < 0.001; ****p < 0.0001. **C:** Comparative SRS spectra from MTZ-treated zebrafish. Orange: control tissue (e.g., liver). Blue: PS-NP–rich glomerular region. Green dashed trace: reference SRS spectrum of 50 nm PS-NPs. PS-characteristic peaks appear at 1008 and 1605 cm^−1^ (fingerprint region) and 3068 cm^−1^ (CH region). **D:** Combined immunofluorescence and SRS imaging of *Tg(pod: NTR-mCherry)* zebrafish cross sections at 5 dpf treated with green fluorescent PS-NPs (-MTZ +PS-NPs), treated with green fluorescent PS-NPs and MTZ (80 µM) (+MTZ + PS-NPs), and without any treatment (-MTZ – PS-NPs). Note that some podocyte signal is still present after treatment with 80 µM MTZ. Scale bar: 50 µm. **E:** Combined immunofluorescence and SRS imaging of *Tg (pod: NTR-mCherry)* zebrafish cross sections at 5 dpf treated with green fluorescent PS-NPs (-MTZ +PS-NPs), treated with green fluorescent 50 nm PS-NPs and MTZ (1 mM) (+MTZ +PS-NPs), and without any treatment (-MTZ -PS-NPs). Podocytes are labeled in red, and PS-NPs are labeled in green. Images were taken with 40X magnification. Scale bar 10 µm. Not that 1 mM MTZ treatment leads to almost complete loss of podocytes (white box) due to cell apoptosis.

In order to determine whether detected signals in MTZ-injured larvae are corresponded to PS-NPs, we acquired SRS spectra from both the glomerular region of MTZ-treated larvae and reference PS-NPs **(Fig. 4C)**. The NP-rich glomerular region exhibited the characteristic PS peaks at 1008 and 1605 cm^−1^ in the fingerprint region and at 3068 cm^−1^ in the CH region, validating the chemical identity of the detected particles.

Combined immunofluorescence and SRS imaging of Tg(pod:NTR-mCherry) zebrafish revealed that residual podocyte fluorescence remained detectable after 24 h of 80 µM MTZ treatment. Subsequent PS-NP exposure resulted in colocalization of green-labeled PS-NPs with the glomerular area **(Fig. 4D)**, suggesting that even moderate podocyte injury is sufficient to allow nanoparticle translocation across the glomerular basement membrane.

To examine the effect of glomerular barrier integrity on NP localization in more detail, we performed combined immunofluorescence and SRS imaging on cross sections of Tg(pod:NTR-mCherry) zebrafish larvae with complete podocyte depletion induced by higher concentrations of MTZ (1 mM). In untreated controls (-MTZ -PS), podocytes were clearly labeled in red, and no PS-NPs were detected (green signal). Treatment with PS-NPs alone (-MTZ +PS-NPs) resulted in the presence of PS-NPs predominantly within the glomerular capillaries, while podocytes remained intact. In contrast, larvae subjected to 1 mM MTZ-mediated podocyte ablation (+MTZ +PS-NPs) showed almost complete loss of podocytes (white box), consistent with apoptosis induced by 1 mM MTZ, and PS-NPs were observed not only in the glomerular region but also in areas normally occupied by podocytes (green signal), indicating that barrier disruption facilitates nanoparticle access to accumulate in the glomerulus **(Fig. 4E)**.

These results reveal that podocyte integrity is a primary determinant of NP exclusion from the glomerulus in the whole-body, living animals, capturing systemic physiology, circulation, and inter-organ interactions that are absent in isolated perfused kidney experiments. It has been shown that even partial podocyte loss permits PS-NPs translocation into normally protected glomerular compartments, establishing a direct link between structural barrier damage and renal nanoparticle accumulation. Furthermore, the integration of SRS spectral analysis with fluorescence imaging provides chemical confirmation of PS-NPs identity *in vivo*, offering mechanistic insight into how glomerular barrier breakdown governs renal nanoparticle handling.

### Accumulation of *PS-NPs in damaged kidney in isolated perfused kidney model*

Following the water exposure of 50 nm PS-NPs, the NPs were detected in the gut but not in the kidney of zebrafish larvae, it was unclear whether this reflects a true exclusion by the renal filtration barrier or limited systemic access due to intestinal uptake and larval vascular immaturity. To address this question, we assessed the renal uptake of PS-NPs under the direct systemic exposure of PSNPs in a mammalian model. To that aim, we applied green fluorescent PS-NPs directly to the isolated perfused kidneys, allowing the assessment of particle–kidney interactions in a controlled setting, independent of gut barrier. In intact kidneys, 50 nm PS-NPs were too large to pass through the glomerular filtration barrier, resulting in their detection within the glomerular capillaries (**Fig. 5A, B** red boxes), but their absence from podocytes and the tubular lumen. To test whether glomerular injury facilitates particle passage, kidneys were pretreated in the perfused state with puromycin aminonucleoside (PAN). PAN treatment is a well-established model to cause glomerular damage in zebrafish and mice models ^43^. Two hours of PAN perfusion induced early podocyte injury, including partial podocyte effacement (**Fig. 5B**, black arrow), as well as acute glomerular endothelial stress with focal areas of endothelial cell loss from the glomerular basement membrane (**Fig. 5B**, red arrow). Following PAN treatment, PS-NPs were detected not only in the glomeruli but also, to a lesser extent, in tubular regions (**Fig. 5A**, white arrow). These results confirm that an intact glomerular filtration barrier effectively prevents renal uptake of 50 nm PS-NPs, whereas glomerular injury compromises barrier selectivity, enabling nanoparticle translocation into glomeruli and downstream tubular regions. This highlights glomerular integrity as a critical determinant of renal nanoparticle exposure and establishes a mechanistic link between barrier disruption and particle accumulation in kidney tissue.

**Figure 5:**
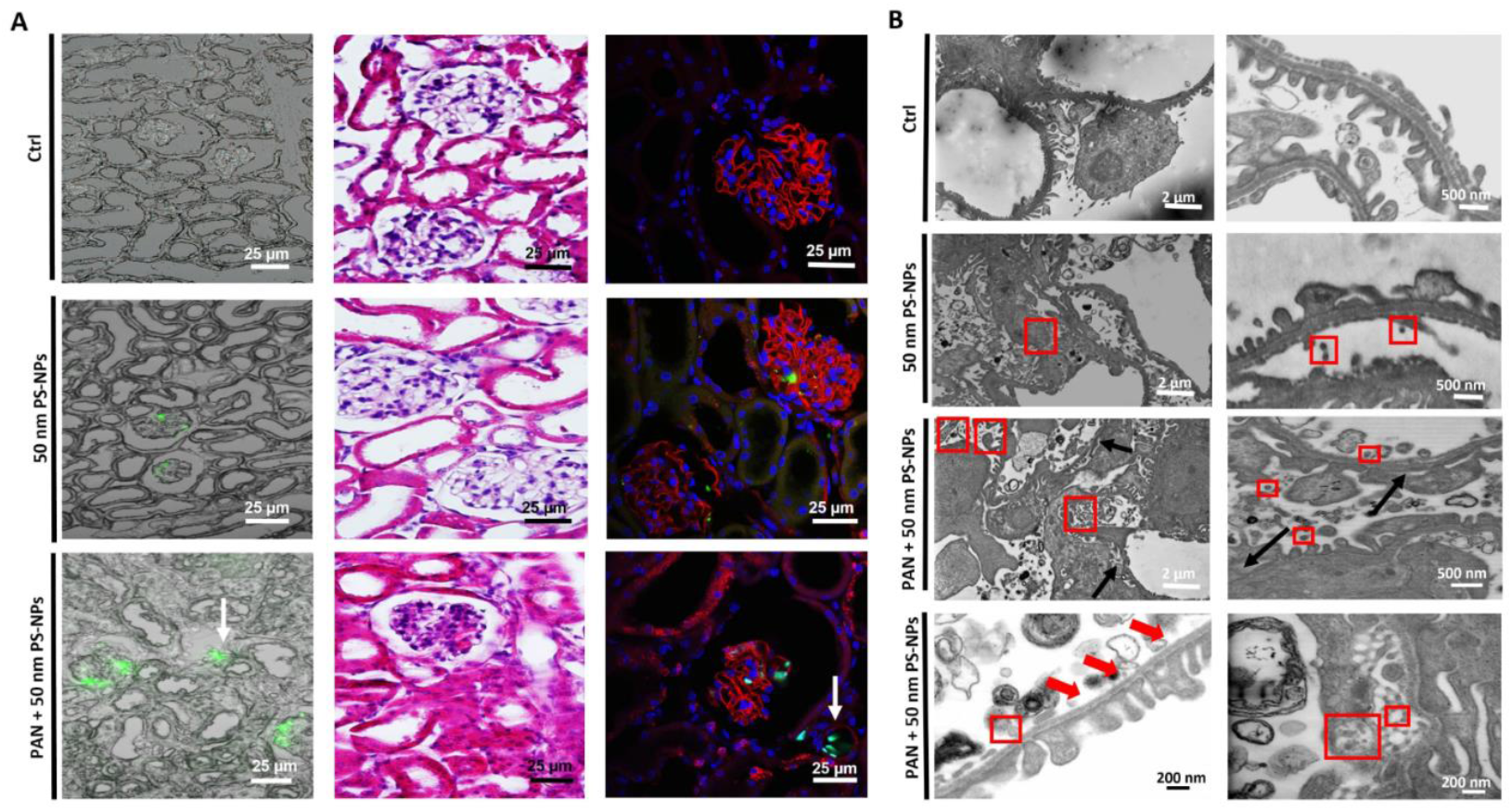
PS-NPs are restricted to pass the glomerular basement membrane in healthy condition. A: Merged images of bright field and fluorescent channel, H&E staining, and immunofluorescent images: nephronectin assigned as red color, DAPI staining assigned the nuclei as blue in the kidney sections of isolated perfused mice kidney. Isolated perfused kidneys were perfused with regular perfusate (Ctrl) versus green fluorescent PS-NPs (PS-NPs) or PS-NPs following puromycin (PAN) treatment for 2h (PAN + PS-NPs). White arrow illustrates green fluorescence of PS-NPS in the tubular region after PAN induced glomerular injury. Scale bar 25 µm. B: Transmission electron microscopy of kidney sections of isolated perfused mice kidney. Isolated perfused kidneys were perfused with regular perfusate (Ctrl) versus green fluorescent PS-NPs (PS-NPs) or PS-NPs following puromycin treatment (PAN) for 2 h (PAN +PS-NPs). In intact kidneys, nanoparticles were accumulated within glomerular capillaries (red boxes) but were absent from podocytes and the tubular lumen. Pretreatment with PAN for 2 h induced early podocyte injury with partial effacement (black arrow) and acute glomerular endothelial stress with focal endothelial cell loss from the glomerular basement membrane (red arrow). Following PAN-induced glomerular damage, PS-NPs were detected within the glomeruli and, to a lesser extent, in tubular regions (white arrow), demonstrating that barrier disruption facilitates nanoparticle translocation. Scale bar 500 nm

### Label-free SRS imaging reveals tissue-specific localization and lipid co-association of PS-NPs after ex vivo glomerular injury

SRS microscopy enabled label-free visualization of PS-NPs distribution within mouse tissue sections and provided chemical contrast information as a complementary method to fluorescence detection. In control tissues, no PS-specific Raman signal was detected, whereas sections from PS-NP-exposed animals displayed distinct signals at the characteristic PS vibrational bands (1008, 1605, and 3068 cm^−1^), co-localizing with the fluorescence signal of the particles (**Fig. 6A, supplementary Fig. 1**). Merged SRS and fluorescence images confirmed that PS-NPs accumulated in glomerular vasculature, often in close spatial proximity to lipid-rich domains visualized through the CH_2_ stretching band at 2850 cm^−1^. Co-localization of PS and lipid channels (white in merged panels of **Fig. 6A supplementary Fig. 1**) indicated intimate particle–lipid interactions, suggesting incorporation or adsorption of nanoparticles within glomerular vasculature. The SRS spectra extracted from particle-rich regions showed clear spectral signals from surrounding tissue, confirming the chemical specificity of PS detection (**Fig. 6B**). Co-exposure to the nephrotoxicant PAN resulted in an altered distribution pattern, with more diffused PS signals and enhanced lipid co-localization, indicating compromised tissue barrier integrity and increased nanoparticle-membrane interactions (**Fig. 6A, B**). These findings demonstrate that SRS provides a powerful, label-free approach to detect NP localization, tissue integration, and biochemical context within complex biological microenvironments. Under physiological conditions, 50 nm PS-NPs were retained in the glomerular capillaries and cannot cross the intact filtration barrier. However, glomerular injury facilitates partial translocation of the particles into podocyte and tubular compartments, confirming that barrier integrity is a critical determinant of renal nanoparticle uptake.

**Figure 6:**
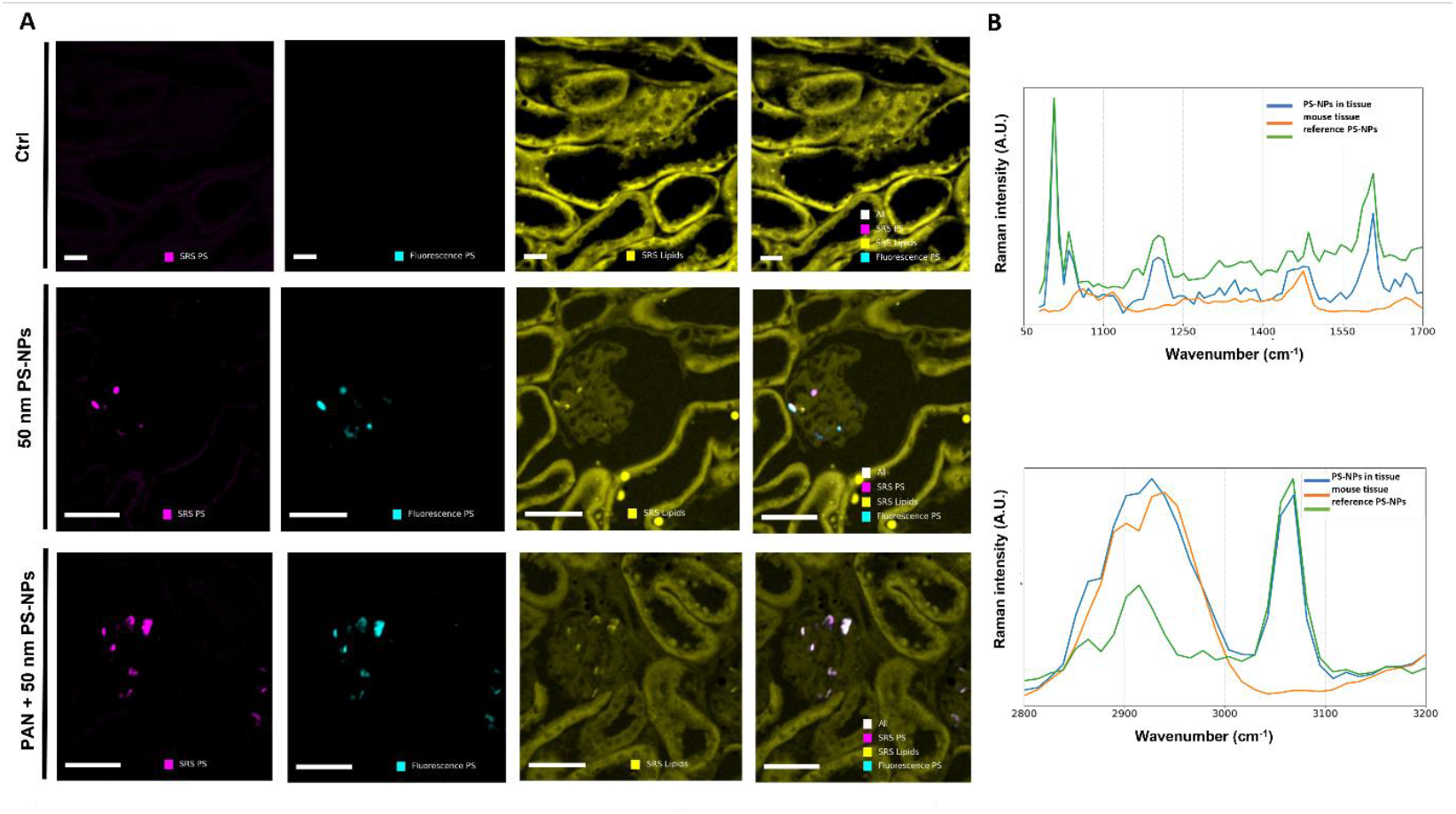
PS-NPs accumulate in glomerular regions after injury ex vivo. A: Representative stimulated Raman scattering (SRS) and fluorescence images showing the spatial localization of green fluorescent PS-NPs and their interaction with the mouse tissue sections. Each group (CTRL, PS-NPs, and PS-NPs +PAN) includes image panels displaying the SRS PS channel (magenta, PS nanoparticle signal), fluorescence PS channel (cyan, fluorescence signal from PS particles), SRS lipids channel (yellow, lipid distribution), and the merged image of all three channels, where white indicates overlapping regions of PS and lipid signals. Scale bar = 20 µm. More figures can be found in supplementary Fig. 1. B: The accompanying SRS spectra show the fingerprint and CH stretching regions, where the green line represents the reference PS-NPs spectrum, the orange line corresponds to the mouse tissue spectrum, and the blue line indicates the PS particle-rich region (ROI) within the mouse tissue. These images and spectra collectively demonstrate the localization of PS nanoparticles within tissue microenvironments, the co-localization with lipid-rich domains, and the spectral distinction between PS particles and endogenous tissue components. Characteristic Raman peaks of PS at 1008, 1605, and 3068 cm^−1^ were detected in PAN-damaged kidneys perfused with 50 nm PS-NPs (upper graph). In contrast, tissue regions displayed dominant signals at 1470-1475 cm^−1^ (CH_2_/CH_3_ bending) and 1667 cm^−1^ (C=C stretching/amide I), allowing spatial distinction of PS-NPs from the surrounding lipid- and protein-rich kidney matrix (lower graph).

### Stimulated Raman spectroscopy detects PS-NP uptake in glomerular cells in vitro

To complement our *in vivo* studies and to directly assess whether glomerular cells can internalize 50 nm green fluorescent PS-NPs in the absence of systemic and barrier constraints, we performed *in vitro* experiments using human glomerular endothelial cells (GECs) and podocytes. This enabled us to study direct effects of high concentrations of PS-NPs on glomerular cells. Exposure of these cells to fluorescently labeled 50 nm PS-NPs demonstrated efficient NP internalization by both cell types. 3D constructed models and lysotracker co-staining confirmed the colocalization of PS-NPs with lysosomes, indicating endolysosomal trafficking in GECs (**Fig. 7A, B**) and podocytes (**Fig. 7C, D**). In addition to fluorescence-based detection, SRS imaging further confirmed PS-NP identification in glomerular cells. In the SRS data **(Fig. 7E)**, the merged image displays the 1008 cm^−1^ PS-NP signal in green and the 1667 cm^−1^ tissue CH/amide signal in yellow, allowing clear visualization of cellular structure and nanoparticle-associated contrast. The corresponding spectra covering both the fingerprint region and the CH region show a strong 1008 cm^−1^ peak for intracellular PS-NPs (blue trace), while the surrounding cell body exhibits only tissue-specific Raman features (orange trace).

**Figure 7:**
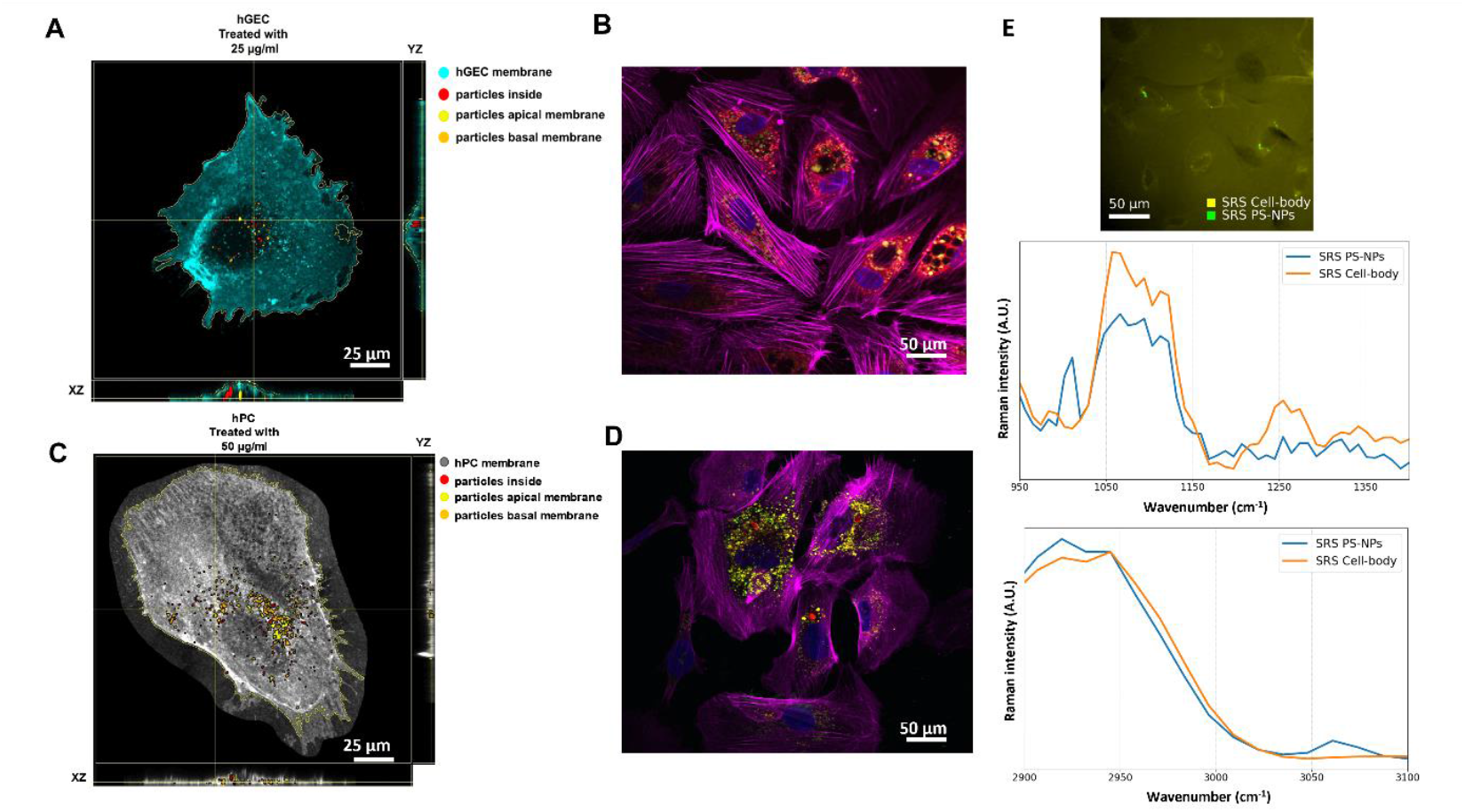
PS-NPs are taken up by glomerular cells. A and C: 3D reconstruction of immunofluorescent images of (A) human glomerular endothelial cells (GECs) and human podocytes (hPCs) (C) with the use of Particle-in-cell-3D model shows the incorporation of green fluorescent PS-NP particles into cells. Particle at apical membrane (yellow), particle inside the cells (red), particles at basal membrane (orange). Scale bar is 25 µm. B and D: Fluorescent imaging of PS-NP uptake in GECs (B) and podocytes (D) using double staining with lysotracker and phalloidin. Lysosomes, NPs, Actin filaments, and nuclei are shown in red, green, magenta, and blue, respectively. The co-localization of NPs and lysosomes are shown as yellow spots. Images were recorded with 63x magnification. Scale bars are 25 µm. E: Merged SRS image of the glomerular region showing the PS-NP–specific signal at 1008 cm^−1^ (green) and the tissue related CH/amide signal at 1667 cm^−1^ (yellow). The pseudo-colored composite highlights tissue structure (yellow) confirms the absence of PS-NP signal in control samples (green). This part also includes the corresponding SRS spectra from the same region: the fingerprint-region spectra (including the 1008 cm^−1^ PS-NP peak), and the CH-region spectra. Scale bar: 50 µm.

Despite robust cellular uptake, PS-NP exposure did not affect the mRNA expression of key glomerular markers in either cell type (**supplementary Fig. 2**), and no obvious changes in actin cytoskeleton of podocytes were observed (**supplementary Fig. 3**). These findings suggest that short-term exposure to 50 nm PS-NPs does not impair fundamental cell identity, indicating that these particles appear relatively inert in standard kidney cell assays. These data are in consistent with previous observations in gut- and liver-derived human cell lines ^46^.

### PS-NPs alter gut microbiota composition in zebrafish

PS-based NPs have previously been shown to accumulate in the intestine and trigger oxidative stress and inflammatory responses ^47^. Similarly, in our study, the gastrointestinal tract represented the primary site of PS-NP accumulation. based on the pronounced intestinal localization and the growing recognition of the gut as a key interface between environmental exposures and systemic physiology ^48^, we examined whether PS-NP exposure alters the gut microbiota composition in zebrafish larvae. Microbiota profiling of PS-NP-exposed zebrafish revealed marked shifts in microbial community composition. Taxonomic analysis at the genus and family levels demonstrated altered relative abundances (**Fig. 8A**). While α-diversity indices (Observed richness, Fisher’s, Shannon, and inverse Simpson indices) showed only a trend toward reduced species richness (**Fig. 8B**) β-diversity analyses (Bray–Curtis dissimilarity and Jaccard distance) revealed significant community divergence between control and PS-NP-treated groups (**Fig. 8C**), confirming compositional restructuring of the microbiome. This pattern of community compositional change without consistent, large α-diversity loss has been observed previously for polystyrene particles in zebrafish and other models and is frequently accompanied by shifts in host metabolism and inflammation ^49^. Linear Discriminant Analysis Effect Size (LEfSe), a statistical approach that identifies bacterial taxa with significantly different relative abundances between experimental groups and estimates their discriminatory relevance, revealed enrichment of *Unibacterium, Chitinimonas, Legionella, Sediminibacterium, Rikenellaceae*, and *Acinetobacter* in control larvae, whereas *Bradyrhizobium* and *Pseudomonas* were enriched following PS-NP exposure (**Fig. 8D**). other studies showed that NPs alter the composition of microbial community, even at doses that do not strongly affect the viability of intestinal epithelial cell viability ^50^. This pattern suggests a shift toward stress-tolerant or inflammation-associated taxa at the expense of commensal bacterial populations.

**Figure 8:**
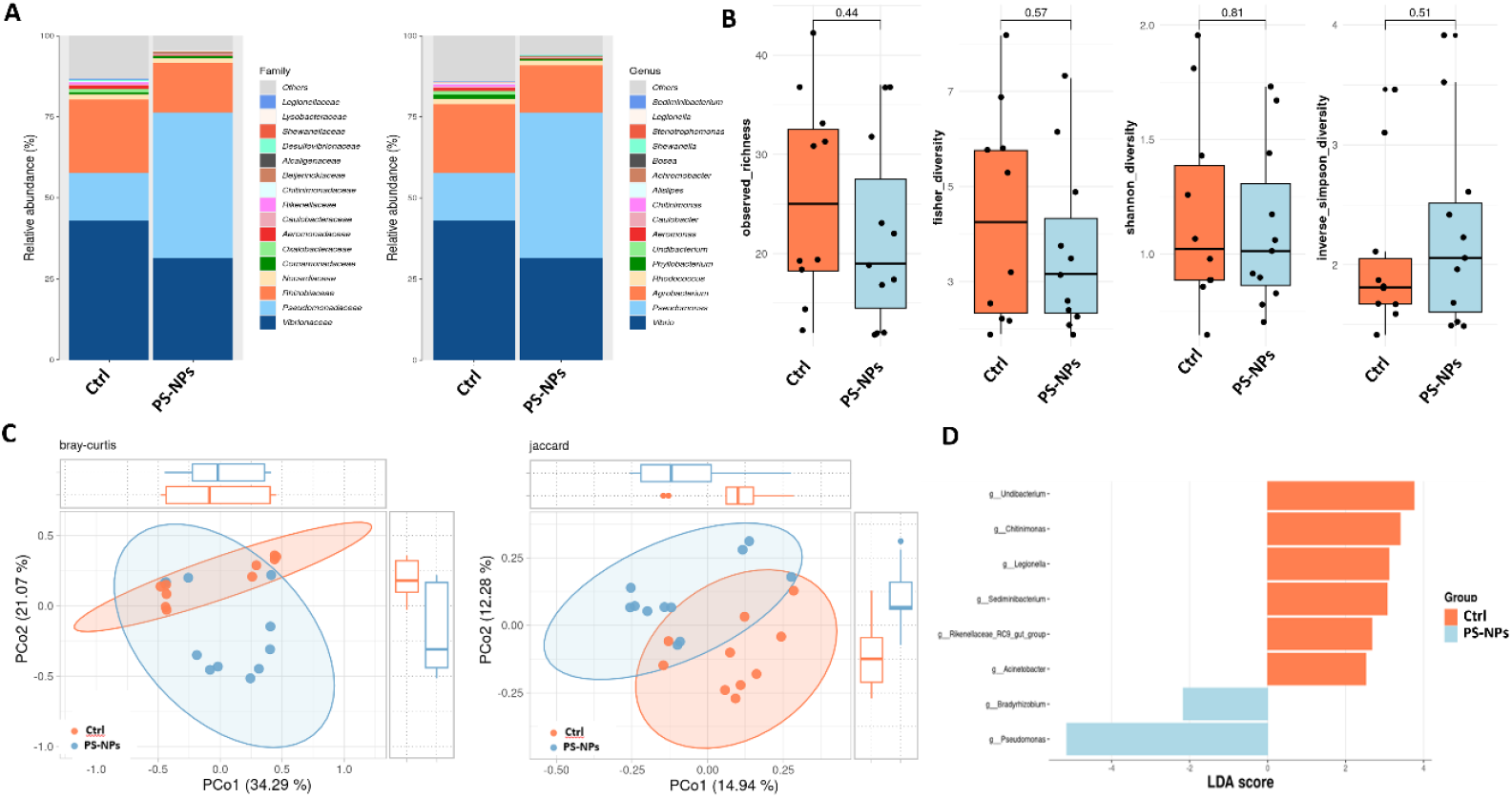
PS-NPs cause changes in zebrafish microbiota. A: Stacked bar plotting shows relative abundances (%) of different bacterial taxa of zebrafish larvae between PS-NP treated groups and controls at genus and family level. n = 10 control (Ctrl) zebrafish larvae and 11 larvae treated with PS-NPs B: Alpha diversity metrics (observed richness, Fisher’s alpha, Shannon diversity index and Simpson index) at ASV level were compared between zebrafish larvae treated with PS-NPs and control zebrafish. Wilcoxon rank-sum tests were used for statistical analysis. C: Beta diversity analysis comparing the (dis)similarity of the microbial composition of controls to PS-NP treated zebrafish larvae using principal coordinates analysis based on Bray curtis dissimilarity and jaccard distance. PERMANOVA tests for significant differences in community composition between groups showed a significant difference between PS-treated and control samples. P<0.001. D: LEfSe analysis (Linear Discriminant Analysis Effect Size) to identify genus that are significantly different between PS-NP treated and control zebrafish larvae.

### PS-NPs trigger host metabolic adaptation

Gut dysbiosis is paralleled by transcriptional signatures of metabolic and stress adaptation in exposed larvae. Bulk RNA sequencing of whole zebrafish larvae exposed to PS-NPs revealed transcriptional upregulation of genes linked to immune and stress responses (*irg1l, mpeg1, cd68, rgs2, prkcd*), xenobiotic metabolism (*cyp8b1, sult5a1*), and epithelial barrier function (**Fig. 9A**). Similar metabolite changes have been shown before ^51^. KEGG analysis revealed that PS-NP exposure caused widespread metabolic disruption in zebrafish larvae, with particular impact in developmental pathways, cofactor biosynthesis, and xenobiotic detoxification (**Fig. 9B**). The pattern suggests systemic toxicity affecting multiple organ systems, with potential implications for energy metabolism, and endocrine function. This coordinated response indicates that PS-NPs modulate a gut–microbiota–host axis, linking microbial shifts to host metabolic adaptation. Expansion of opportunistic bacteria such as *Pseudomonas* may contribute to pro-inflammatory signaling, while host metabolic reprogramming likely represents an adaptive mechanism to NP-induced stress.

**Figure 9:**
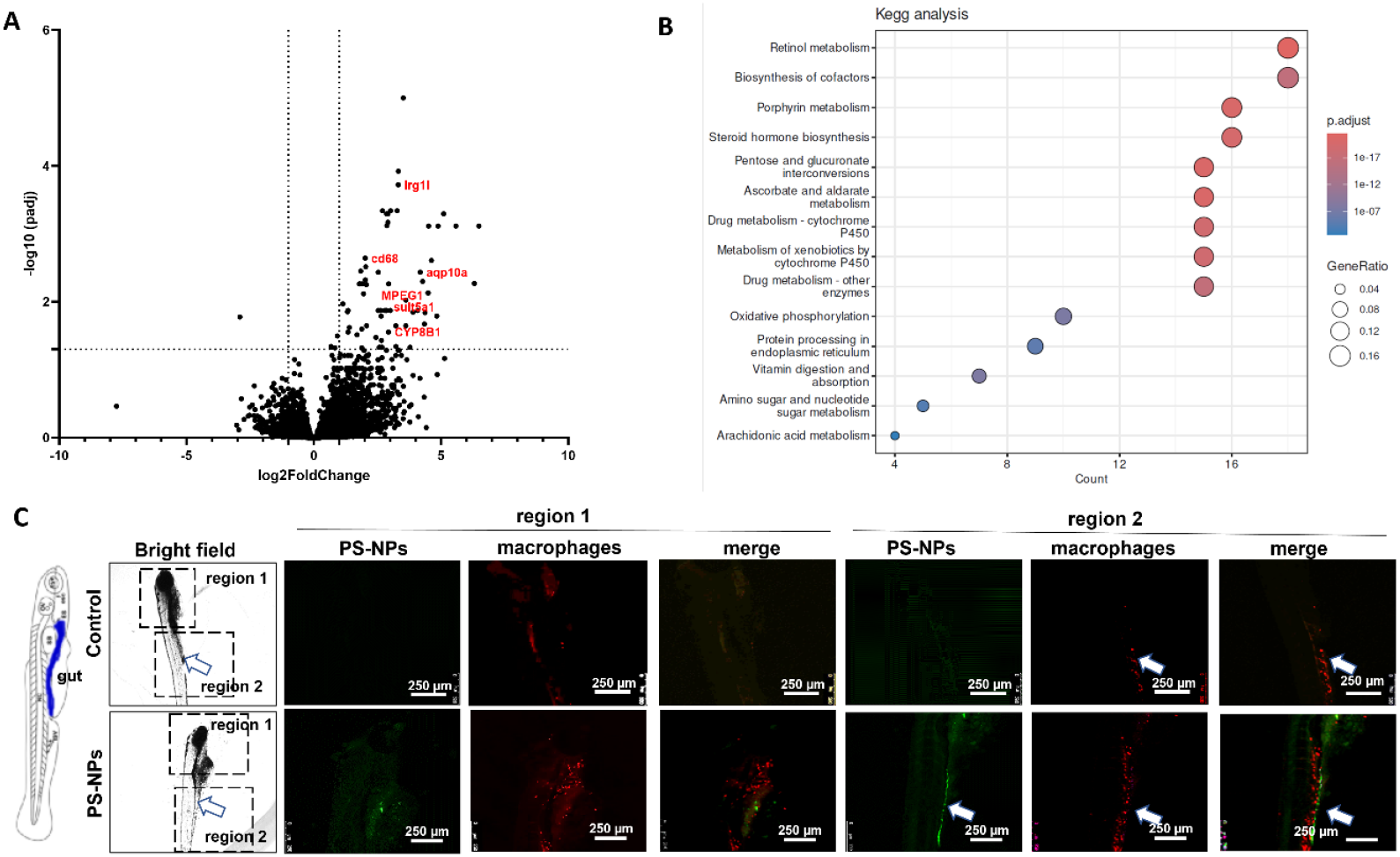
Upregulation of inflammatory marker and recruitment of myeloid immune cells to the intestine following PS-NP exposure. A: Volcano-Plot of bulk RNA sequencing data showing log2fold change and -log10 (p-values) of expression changes in zebrafish mRNA from PS-NPs exposed zebrafish larvae compared to Ctrl. n = 10 control (Ctrl) zebrafish larvae versus 11 larvae treated with PS-NPs. B: Kegg pathway analysis of bulk RNA sequencing data showing genes regulated in zebrafish larvae after treatment with PS-NPs. n = 10 control (Ctrl) zebrafish larvae versus 11 larvae treated with PS-NPs C: Bright field and fluorescent images of whole transgenic zebrafish larvae expressing a red fluorescent reporter under the control of the *lysozyme C* (*lysC*) promoter, labeling a subset of myeloid cells (including macrophages and some granulocytes). Zebrafish were exposed to PS-NPs within the fish water or left untreated (Ctrl). Following treatment, a pronounced accumulation of fluorescent myeloid cells was observed in the intestinal region (arrowhead), indicating localized immune cell recruitment.

Therefore, PS-NP exposure reshaped the gut microbiota, promoting stress-tolerant and inflammation-associated taxa, while simultaneously triggering a coordinated host response that involved immune activation, xenobiotic metabolism, and epithelial barrier function. This highlights the systemic impact of nanoparticle-induced changes in the gut. PS-based NPs have been shown to accumulate in the intestine, causing oxidative stress and inflammatory responses despite of the weak acute *in vitro* toxicity ^11, 16^.

### PS-NPs recruit immune cells to the gut

To investigate the recruitment of myeloid immune cells upon exposure to green fluorescent PS-NPs, we utilized a transgenic zebrafish line expressing a red fluorescent reporter under the control of the lysozyme C promoter (*Tg(lyz:DsRed2)*, which labels a subset of myeloid cells, including macrophages and some granulocyte ^29^. Following PS-NP treatment, we observed a pronounced accumulation of fluorescent immune cells in the intestinal region (**Fig. 9C**, arrowhead), indicating a localized immune response. These findings suggest that PS-NPs trigger site-specific recruitment of myeloid cells, highlighting the gut as a primary interface for nanoparticle-induced immune activation *in vivo*. If the accumulation of immune cells was a direct effect of the particles or mediated by microbiota changes seen after NP exposures require manipulative experiments with antibiotic depletion, germ-free or gnotobiotic zebrafish along with validation of key taxa/metabolites using orthogonal assays. Experimental evidence that MP/NPs damage zebrafish kidneys and suppress innate immune pathways supports this model ^52^ and aligns with our findings of edema after PS-NP treatment even though PS-NPs did not accumulate in the kidney

In our study, we employed spherical PS-NPs with a nominal diameter of 50 nm as a model NP to investigate particle–host interactions. Although NPs in the environment exhibit heterogeneous morphologies, spherical PS-NPs represent a relevant and well-characterized proxy for several reasons. First, at the nanoscale, surface tension and polymer chain mobility favor the formation of near-spherical particles, and consequently, spherical NPs are among the most likely naturally occurring shapes following progressive fragmentation and weathering of larger plastics ^53^. Second, using monodisperse spherical NPs enables controlled assessment of size-dependent biological effects, independent of shape-related confounders such as fiber length or aspect ratio. Finally, PS-NPs are well-established reference materials in nanotoxicology and serve as reproducible surrogates to explore mechanisms of uptake, biodistribution, and toxicity. Thus, spherical 50 nm PS-NPs provide a reproducible and mechanistically informative model system to study NP-host-interactions under controlled experimental conditions. However, findings obtained using such model particles should be understood as establishing upper-bound and principle-based barrier for interactions rather than serving as direct representations of environmentally aged NPs. This approach provides mechanistic framework to guide future studies that incorporate particle heterogeneity and environmental aging.

In conclusion, our results support a model in which PS-NPs mainly influence the gut, altering microbial communities, initiating local immune responses, and triggering systemic metabolic changes. Renal accumulation occurs only when there is compromise in the glomerular barrier. Importantly, the edema observed in zebrafish larvae likely results from the combination of factors including: disruption in the gut barrier, systemic metabolic and inflammatory responses, and potential contributions from non-renal osmoregulatory tissues, such as skin. These findings highlight the importance of considering multi-organ interactions when interpreting systemic responses to environmental nanoparticles. Our study emphasize the robustness of the glomerular filtration barrier as a conserved mechanism that restrict nanoparticle entry under physiological conditions. By combining zebrafish *in vivo* models, isolated perfused kidneys, and *in vitro* cell assays with SRS imaging, we provide a multi-scale mechanistic framework that enhance our understanding of the broader ecological and human health risks associated with NPs.

## Abbreviations

DLS: dynamic light scattering
Dpf: day post fertilization
LEfSe: linear discriminant analysis effect size
MPs: microparticle
MTZ: metronidazole
NPs: nanoparticle
PAM: puromycin aminonucleoside
PBS: phosphate-Buffered Saline
PFA: paraformaldehyde
PS-NP: polystyrene nanoparticle
SRS: stimulated Raman scattering
SEM: scanning electron microscopy
TEM: transmission electron microscopy

## CRediT authorship contribution statement

**MY:** Writing – review & editing, Methodology, Data curation.

**MK:** Methodology, Data curation, Writing – review & editing

**GS:** Methodology, Data curation, Writing – review & editing

**SW:** Methodology, Data curation, Writing – review & editing

**AO:** Methodology, Data curation, Writing – review & editing

**FS:** Methodology, Data curation, Writing – review & editing.

**MS:** Resources, Funding acquisition, Writing – review & editing.

**SC:** Resources, Funding acquisition, Writing – review & editing.

**JMD:** Writing – review & editing, Writing – original draft, Supervision, Resources, Project administration, Funding acquisition.

## Acknowledgements

The excellent technical assistance of Robert Götz is deeply acknowledged.

