## Supplementary Figure for "Gut and Glomerular Barriers Determine Nanoplastic Fate and Systemic Impact"

***Shared first authorship**

**#Corresponding author**

Janina Müller-Deile, Department of Nephrology and Hypertension, Uniklinikum Erlangen, Friedrich-Alexander-Universität (FAU) Erlangen-Nürnberg, Erlangen, Germany,

### Supplements

### Table of content

Supplementary Figure 1

Supplementary Figure 2

Supplementary Figure 3

Supplementary table 1

**Supplementary Figure 1:** ***PS-NPs accumulate in glomerular regions after injury ex vivo.***

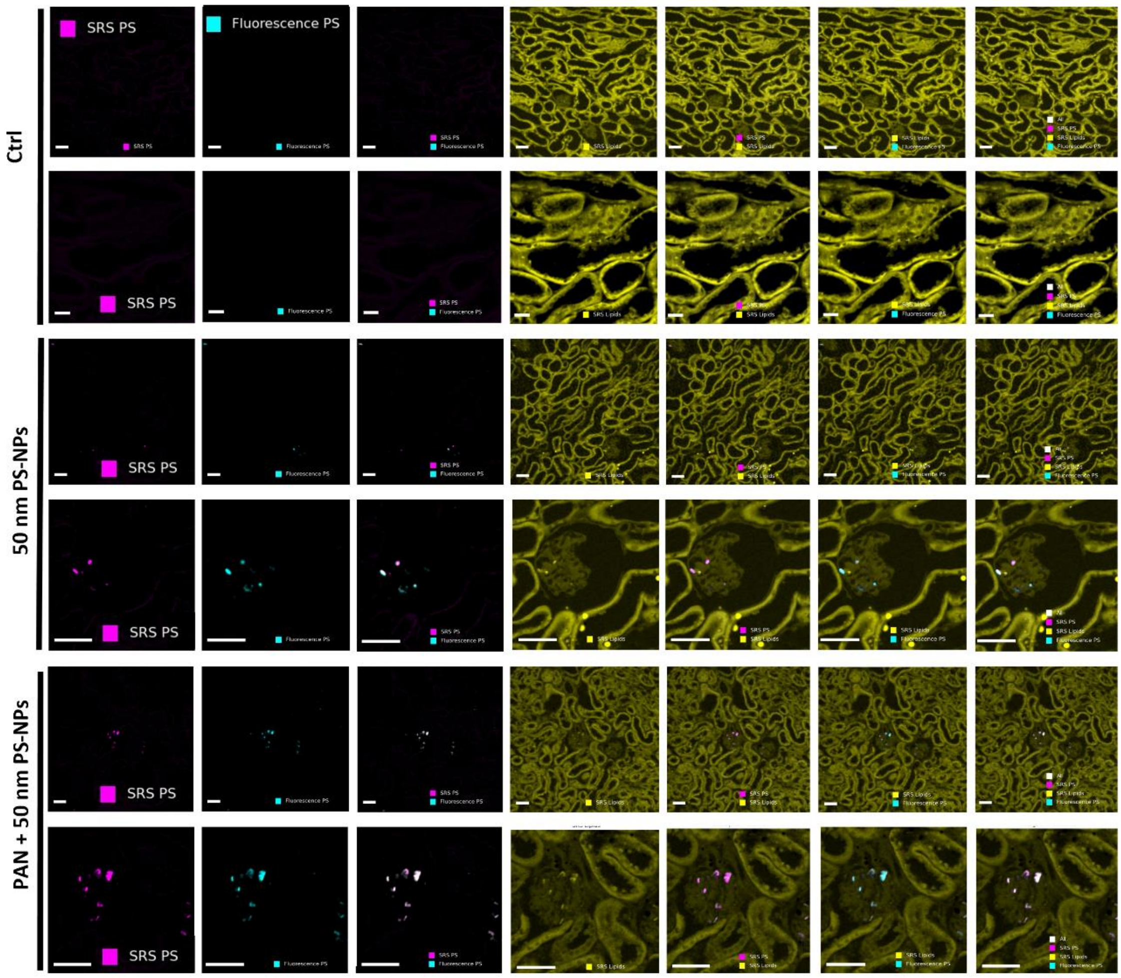


Representative stimulated Raman scattering (SRS) and fluorescence images showing the spatial localization of green fluorescent PS-NPs and their interaction with the mouse tissue sections. Each group (CTRL, PS-NPs, and PS-NPs +PAN) includes seven image panels displaying the SRS PS channel (magenta, PS nanoparticle signal), fluorescence PS channel (cyan, fluorescence signal from PS particles), merged SRS PS and fluorescence PS channels, SRS lipids channel (yellow, lipid distribution), merged SRS PS and SRS lipids channels, merged SRS lipids and fluorescence PS channels, and the merged image of all three channels, where white indicates overlapping regions of PS and lipid signals. For each group, the upper row shows the full field of view (scale bar = 50 µm) and the lower row presents a magnified region of interest (ROI) from the same field (scale bar = 20 µm).

**Supplementary Figure 2:** ***Expression of podocyte and glomerular endothelial cell makers following exposure to PS-NPs.***

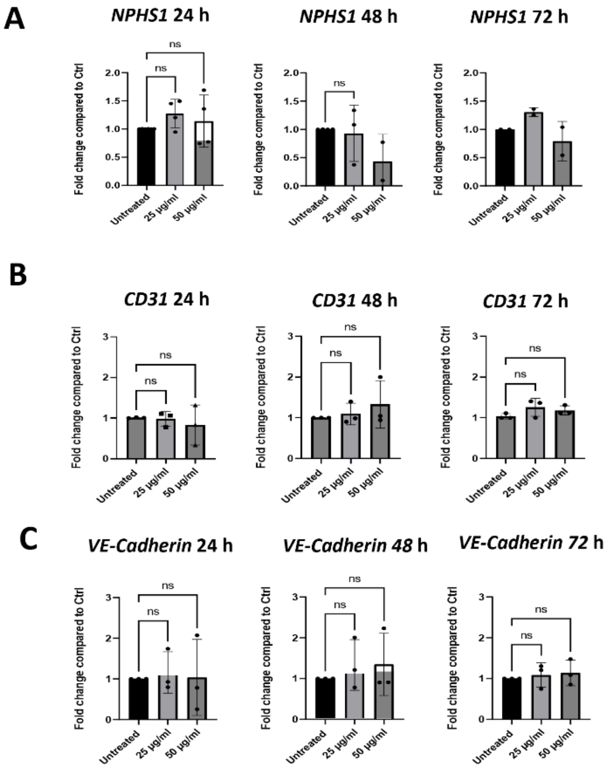


A: mRNA expression of *NPHS1* after exposure of cultured human podocytes to PS-NPs (25 µg/mL, 50 µg/mL) or untreated for 24, 48 or 72 h. n = 3 +/- SEM.

B, C: mRNA expression of *CD31* (B) and *Cadherin* (C) after exposure of cultured human glomerular endothelial cells to PS-NPs (25 µg/mL, 50 µg/mL) versus to untreated ones for 24, 48 or 72 h. n = 3 +/- SEM.

**Supplementary Figure 3:** ***Actin cytoskeleton staining after exposure to PS-NPs to podocytes.***

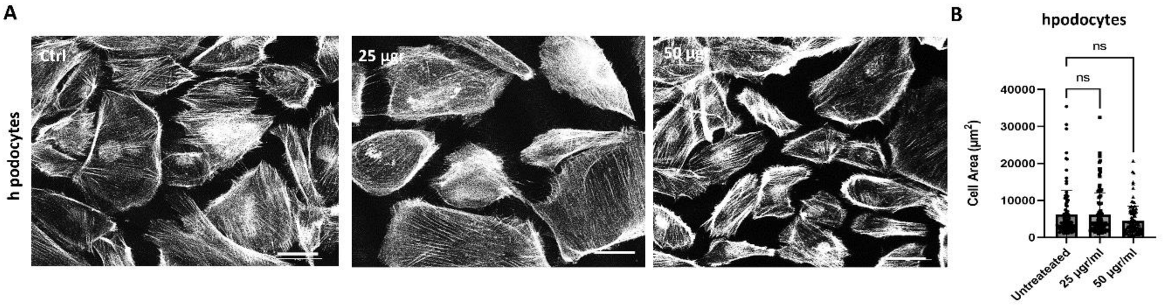


A: Phalloidin staining to visualize actin cytoskeleton of cultured human podocytes in control group (Ctrl) versus treated ones with 25 µg/ or 50 µg PS-NPs. Scale bar = 25 µm.

B: Quantification of cell surface area in µm² of cultured human podocytes in control group (Ctrl) versus treated ones with 25 µg/ or 50 µg PS-NPs.

**Supplementary table 1**

***Primer sequence used in the qPCR experiments.***

CD31 h-PECAM1-fw TGGAAAGCAGATACTCTAGAACGG

CD31 h-PECAM1-rv GGGATGTGCATCTGGCCTT

VE-Cadherin h-CDH5-fw TGGTGGAAGCGCGAGAT

VE-Cadherin h-CDH5-rv AAATGTGTACTTGGTCTGGGT

h-NPHS1-fw AGGACCGAGTCAGGAACGAA

h-NPHS1-rv TCTGTTGTGCTGACCGTGG
